# Collateral sensitivity interactions between antibiotics depend on local abiotic conditions

**DOI:** 10.1101/2020.04.28.065623

**Authors:** Richard C. Allen, Katia R. Pfrunder-Cardozo, Alex R. Hall

## Abstract

Mutations conferring resistance to one antibiotic can increase (cross resistance) or decrease (collateral sensitivity) resistance to others. Drug combinations displaying collateral sensitivity could be used in treatments that slow resistance evolution. However, lab-to-clinic translation requires understanding whether collateral effects are robust across different environmental conditions. Here, we isolated and characterized resistant mutants of *Escherichia coli* using five antibiotics, before measuring collateral effects on resistance to other antibiotics. During both isolation and phenotyping, we varied conditions in ways relevant in nature (pH, temperature, bile). This revealed local abiotic conditions modified expression of resistance against both the antibiotic used during isolation and other antibiotics. Consequently, local conditions influenced collateral sensitivity in two ways: by favouring different sets of mutants (with different collateral sensitivities), and by modifying expression of collateral effects for individual mutants. These results place collateral sensitivity in the context of environmental variation, with important implications for translation to real-world applications.

## Introduction

As a result of antibiotic use, resistance to antibiotics is increasing in bacterial populations^1^, necessitating efforts to identify new antibiotic types^2^. However, the time, money and risks involved in getting new therapeutics to the clinic^3^, and the targeting of many essential bacterial pathways by existing antibiotics mean that the rate of development for new antibiotics is outstripped by rates of resistance development^3^. To tackle the threat of antibiotic resistance, we must investigate strategies to slow the spread of resistance to existing treatments and to any new treatments in development^4^. One strategy that shows promise in slowing the evolution of resistance is to exploit collateral sensitivity interactions^5–8^. These have been observed for specific combinations of antibiotics where mutations conferring resistance to one antibiotic sensitise bacteria to a second antibiotic^5–8^, thereby increasing its effectiveness and reducing the potential for resistance evolution to the second antibiotic^6,9^. For collateral sensitivity interactions to be exploited therapeutically, it is important that their emergence across different populations of bacteria, such as those in different patients or in different communities, is repeatable. That is, unless collateral sensitivity interactions are predictable, exploiting them in new treatment strategies will be very challenging^10–13^.

Recent work revealed important genetic factors influencing the predictability of collateral sensitivity, but the importance of local abiotic conditions is still unclear. For example, high-throughput *in vitro* studies showed different replicate populations exposed to the same antibiotic sometimes acquire collateral sensitivity to another antibiotic, and sometimes do not^10,11^. This can be explained by different mutations, which vary in their phenotypic effects on resistance, spreading in different replicate populations^11,12,14,15^. However, we know from past work that phenotypic effects of antibiotic resistance mechanisms also vary strongly depending on local environmental conditions^16–18^. For example, bile can upregulate efflux pumps^19^, zinc can reduce the activity of aminoglycoside degrading enzymes^20^, and high temperature can modulate the effects of rifampicin-resistance mutations on growth in the absence of antibiotics^21^. This raises the possibility that local environmental conditions could influence both the emergence of collateral sensitivity (by affecting which of the possible pathways to resistance are most strongly selected during antibiotic exposure) and its expression (by modifying the phenotypic effects of resistance alleles when bacteria are exposed to a second antibiotic). To date, research on collateral sensitivity interactions has focused on testing many combinations of antibiotics^5,6,9,14^, multiple strains^10^, or many replicate populations for individual antibiotic combinations^11^. Therefore, the role of local abiotic conditions in the emergence and expression of collateral sensitivity interactions remains unclear. Answering this question would improve our understanding of the robustness of collateral sensitivity across different populations and environments. This would in turn boost our ability to predict pathogen responses to treatment regimens that exploit collateral sensitivity interactions.

To address these gaps in our knowledge, we tested for collateral effects (cross resistance or collateral sensitivity) between five pairs of drugs, each in four different experimental environments. Each drug pair consisted of a selection drug (which we used in mutant isolation) and a paired drug (which we used to test for collateral effects). We chose pairs of drugs indicated by past work to at least sometimes display collateral sensitivity interactions^5,6^. The four experimental environments were (i) basal, nutrient-rich broth (lysogeny broth, LB, at 37°C and buffered at pH 7.0), plus three types of abiotic environmental variation which we expect to be relevant to pathogens in vivo: (ii) reduced pH (pH 6.5), as found in certain body compartments including abscesses and parts of the gastrointestinal (GI) tract^22,23^, (iii) increased temperature (42°C), as found in companion and livestock animals with higher core temperatures than humans^24^, and (iv) the presence of bile salts (0.5g/L bile salts), which bacteria must contend with in the GI tract^25^. We isolated and sequenced resistant mutants after exposure to each selection drug in each experimental environment. We then tested their resistance phenotypes (IC_90_, 90% inhibitory concentration) for the relevant selection drug and paired drug, again across all four experimental environments. Unlike past work, this manipulation of the experimental environment, during both isolation and phenotyping, in a fully factorial design allowed us to quantify the effects of local abiotic conditions on the emergence (which mutations appear in which treatments?) and expression (in which experimental environments do we see collateral effects from particular mutants?) of collateral sensitivity for multiple candidate drug pairs. Although it was not our aim here to investigate the molecular mechanisms by which collateral effects arise, we place our results in the context of available physiological information where possible.

## Results

### Different resistance mutations emerge in different abiotic conditions

We used whole-genome sequencing to identify genetic changes relative to the ancestral *E. coli* strain for 95 resistant mutants (Figure 1), each isolated after exposure to one of five antibiotics (*selection drug*, the antibiotic present on the agar plate used to select resistant mutants from ancestral cells) in four different abiotic environments (*selection environment*, the local abiotic conditions during overnight growth prior to plating and on the agar plate). We found different genes were mutated depending on which antibiotic bacteria were selected against (main effect of selection drug, PermAVOVA: F_4,94_ = 5.46, p<0.001). However, we also found different genes were mutated depending on local abiotic conditions during the experiment (main effect of selection environment: F_3,94_ = 2.48, p<0.001). The effect of selection environment varied among antibiotics (selection drug by selection environment interaction: F_12,94_ = 1.76, p < 0.001). There was an effect of selection environment only for mutants selected against cefuroxime (F_3,17_ = 7.04, p<0.001), streptomycin (F_3,18_ = 2.27, p<0.05), and trimethoprim (F_3,22_ = 1.97, p<0.05). For example, we observed mutations in the penicillin binding protein *ftsI* when we selected for cefuroxime resistance, but only in the presence of bile (Figure 1). Similarly, *folA* mutations were more common when we selected for trimethoprim resistance in basal and low-pH environments (Figure 1), than in bile and high-temperature environments. In summary, the types of resistance mechanisms that emerged during our mutant screen depended on the local abiotic conditions. To investigate whether this effect was linked to variation of resistance phenotypes across resistance mutations and across abiotic environments, we assayed the selected mutants for resistance phenotypes to their corresponding selection drug in all four abiotic conditions.

**Figure 1:**
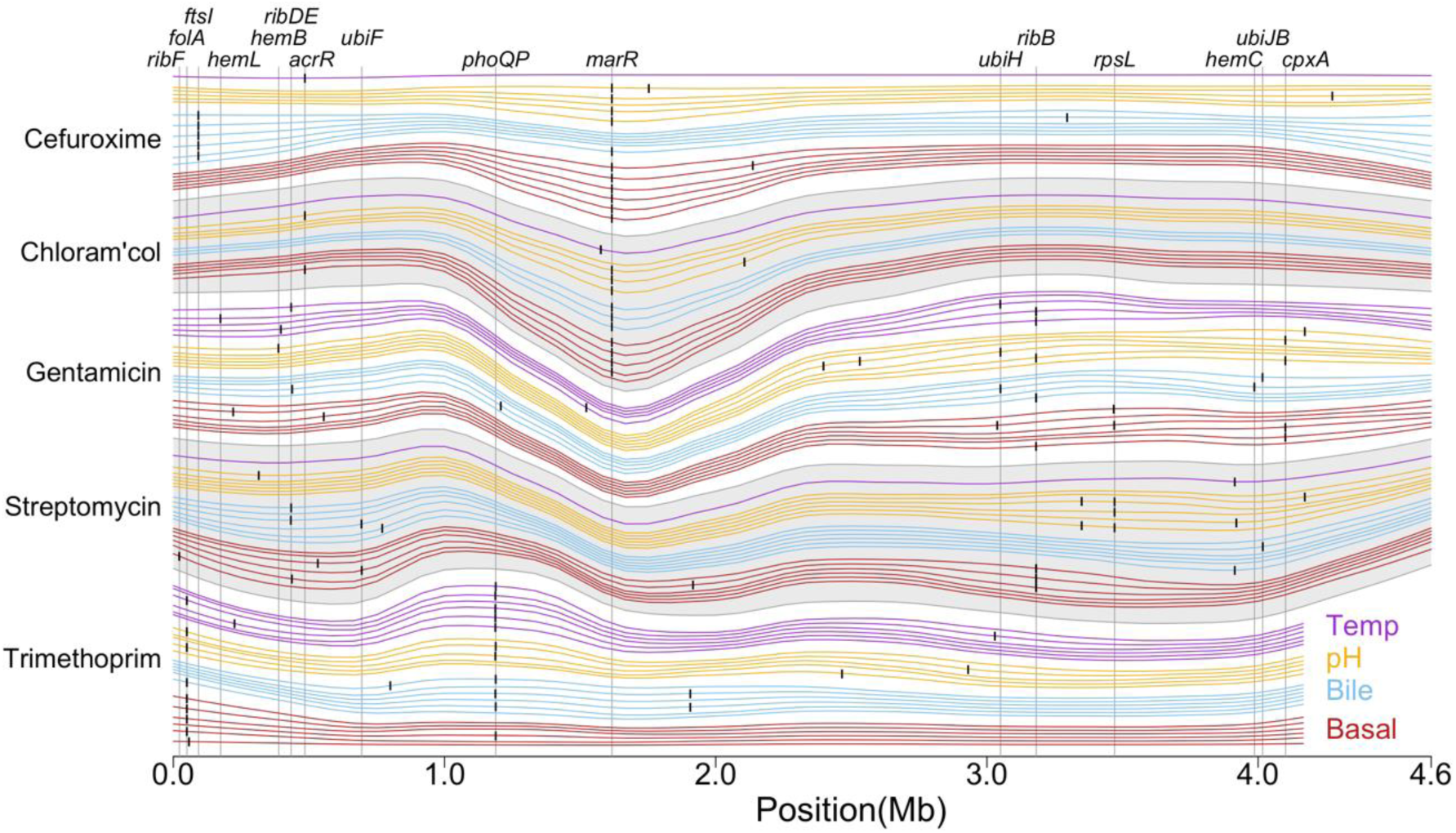
Genotypes of mutants selected for resistance to different antibiotics in different selection environments. Each horizontal line represents a mutant genotype, coloured by the environment where resistance was selected (selection environment, bottom right): basal conditions (red, pH 7.0, 37°C), with the addition of bile (blue, 0.5g/L bile salts), reduced pH (yellow, pH 6.5) or increased temperature (purple, 42°C). Genotypes are grouped according to which drug they were selected against (*y*-axis). Vertical dashes represent mutated regions; these mutated regions are spaced out to better separate the different genotypes (lines) at mutated loci. Gene families that were mutated in three or more independent resistant mutants are labelled at the top; note this includes some gene families where constituent genes are found at multiple separate positions in the genome (*ribFDEB, hemLBC* and *ubiFHJB*). Chloram’col is short for chloramphenicol.

### Resistance to each selection drug depends on which gene is mutated and local abiotic conditions

For all antibiotics, mutants varied in their average resistance against the antibiotic used during isolation (selection drug) depending on which gene family was mutated (effect of genotype on fold-change in IC_90_ relative to the ancestor in the same conditions: Cefuroxime, F_4,19_ = 13.7, p<0.0001; Chloramphenicol F_3,79_ = 36.4, p<0.0001; Gentamicin χ^2^_3_ = 42.4, p<0.0001; Streptomycin, F_6,12_ = 19.9, p<0.0001 and Trimethoprim, χ^2^_4_ = 57.4, p<0.0001; Fig. 2). For example, mutations affecting *ubi* (involved in ubiquinone synthesis) and *rib* (involved in riboflavin synthesis) both conferred increased gentamicin resistance across all assay environments (Fig. 2c), but this effect was stronger for *ubi* mutants. For *ubi* mutants (and possibly *rib* mutants), resistance probably results from alterations to the electron transport chain^26^, reducing membrane potential which aminoglycosides rely on for uptake^27^. Thus, of the observed pathways to resistance in our experiment, some conferred larger increases in resistance than others.

**Figure 2:**
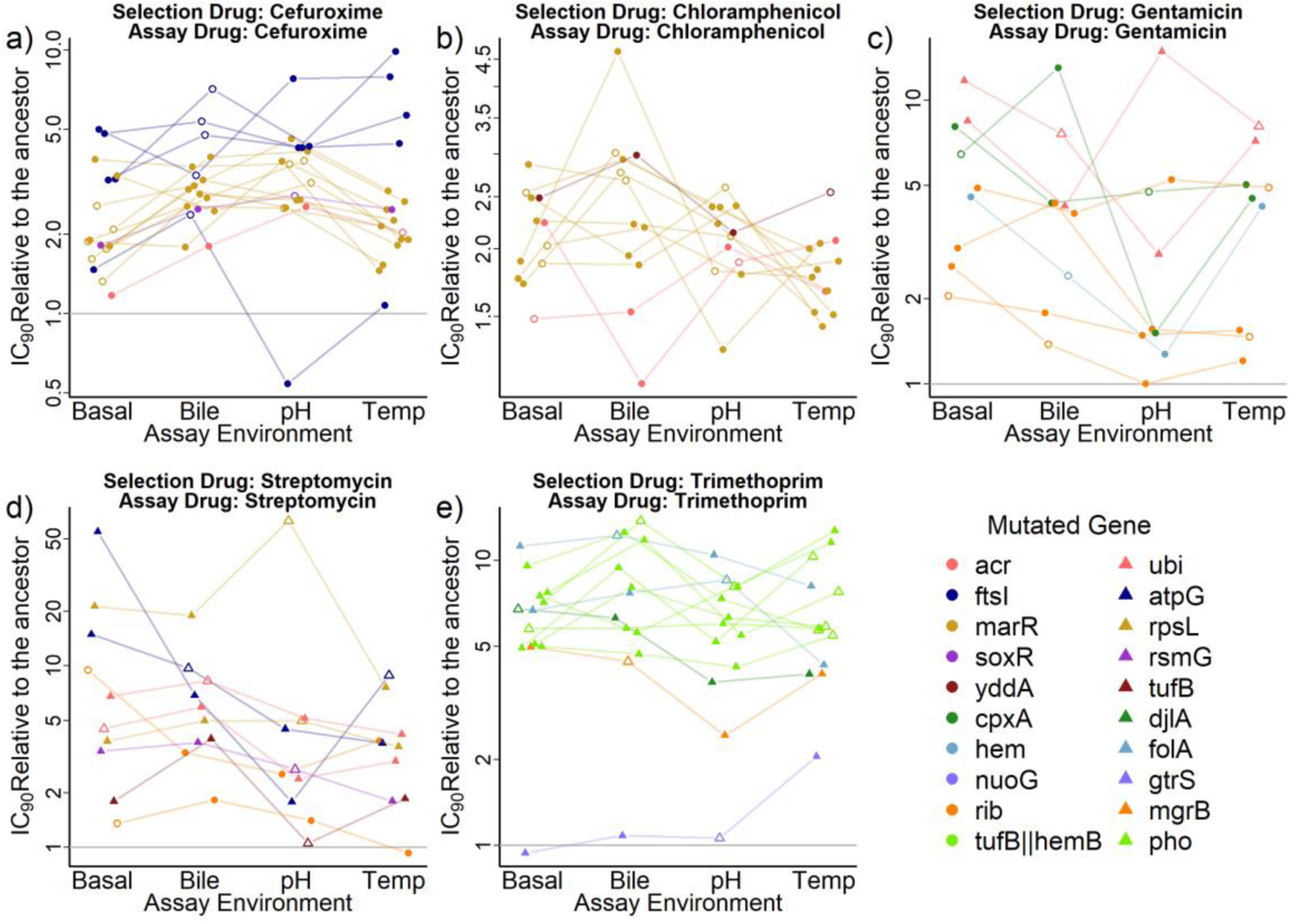
Resistance of each mutant to the antibiotic they were isolated against (*selection drug* = *assay drug*), measured in four sets of abiotic conditions (*assay environment*). Each set of four connected points is a single resistant mutant that was selected for resistance to a) cefuroxime, b) chloramphenicol, c) gentamicin, d) streptomycin or e) trimethoprim. Mutants are coloured according to the gene or gene family that was mutated. Resistance is shown relative to the IC90 of the ancestral strain measured in the same assay environment (IC_90_s for the ancestor across assay environments are shown in Fig. S1). Hollow points indicate sympatric combinations (selection environment = assay environment). Each point is the mean of 2-4 independent replicates (mean *n* = 3.82). The *y* axis is log transformed and the scale varies between panels.

We next asked whether observed variation of resistance relative to the ancestral strain among different genotypes (with mutations in different genes) depended on the *assay environment* (the abiotic conditions during resistance testing). This was the case for mutants selected against cefuroxime (genotype by assay environment interaction: χ^2^_9_ = 21.2, p<0.05), chloramphenicol (χ^2^_6_ = 18.6, p<0.01) and streptomycin (χ^2^_15_ = 33.1, p<0.01), but not gentamicin (χ^2^_9_ = 5.75, p>0.5) and trimethoprim (χ^2^_12_ = 17.1, p>0.1). Some of these effects are consistent with the known physiological functions of affected genes. For example, both *acr* mutants and *mar* mutants had increased resistance to chloramphenicol, but *acr* mutants had relatively weak resistance when assayed in the bile environment (Fig. 2b). This may reflect the different roles of these genes in expression of the acrAB efflux pump (affected by mutations in both *acr* and *mar*^28^) and other protective functions (associated with *mar* mutations only^29^). The smaller change in chloramphenicol resistance relative to the ancestor for *acr* compared to *mar* mutants in the bile treatment may be because acrAB is induced by bile even in the ancestral strain^19^. By contrast, mutations in the *mar* operon maintained their chloramphenicol resistance relative to the ancestor, probably because of the other protective functions regulated by *mar*^29^.

Despite observing that different genes were mutated in different selection environments, and that different mutated genes resulted in variable resistance phenotypes, there was no significant variation of mean resistance among sets of mutants from different selection environments (effect of selection environment on resistance, likelihood ratio test: p>0.05 for all selection drugs). Furthermore, we found no evidence that average resistance was higher for mutants tested in sympatric environments (selection environment = assay environment) than in allopatric environments (selection environment ≠ assay environment) for any of the five selection drugs (Difference between sympatric and allopatric combinations, likelihood ratio test: p>0.05 in all cases). Thus, despite local abiotic conditions influencing which mutants emerged in our screen and how their resistance phenotypes were expressed, this did not result in average differences in resistance among selection environments or a pattern of local adaptation^30,31^ in terms of maximal antibiotic concentrations that mutants could grow in (IC_90_).

We next analysed an alternative measure of resistance, growth of each mutant at the antibiotic concentration used during selection (GASC). Our rationale here was that the mutations most beneficial during our screen (and most likely to result in formation of viable, sampled colonies) are not necessarily the mutations that confer the largest increases in IC_90_. Therefore GASC potentially provides additional information about why some types of mutants were associated with particular selection environments. GASC was calculated from the same dose response curves as the IC_90_ and was positively correlated with IC_90_ across all mutants (correlation: τ=0.61, p<0.0001, Fig. S2). Like IC_90_, variation of GASC was predicted by which gene was mutated and by the interaction between mutated gene and assay environment (p<0.05 for main effect for all antibiotics; p<0.05 for interaction term for 4 antibiotics; Fig. S3). Unlike IC_90_, GASC varied significantly among sets of cefuroxime-resistant mutants selected in different abiotic conditions, with bile-selected mutants performing best (effect of selection environment: χ^2^_3_ = 10.7, p<0.05; Fig. S3a). Thus, addition of bile biased our screen toward a relatively narrow set of mutants that grew well at the cefuroxime concentration used during selection (in particular, *ftsI* mutants; Fig. 1). Consistent with this, we observed resistant colonies in our mutant screen in fewer replicate populations exposed to bile+cefuroxime than other cefuroxime-selection environments (Fig. S4). Note *ftsI* mutants also had relatively high IC_90_s on average (Fig. 2), although this did not translate to significant variation of mean IC_90_ among selection environments as tested above. For trimethoprim-selected mutants, average GASC did not vary among mutants from different selection environments (effect of selection environment: p>0.1). However, there was evidence of local adaptation, in that GASC was higher in sympatric (assay environment = selection environment) than allopatric (assay environment ≠ selection environment) combinations (effect of sympatry: β=0.063, χ^2^_1_ = 8.30, p<0.01). For streptomycin (the other antibiotic where selection environment influenced which genes were mutated), the effect of sympatry was positive but just above the significance cut-off (effect of sympatry on GASC: β=0.04, χ^2^_1_ = 3.81, p=0.051). In summary, analysing GASC supported the variation of resistance depending on mutated gene and assay environment, as in our analyses of IC_90_ above, and revealed additional evidence that some selection environments favoured particular types of resistance mutations.

### Collateral sensitivity and cross resistance depend on resistance mechanism and local abiotic conditions

We next tested whether resistant mutants for each selection drug showed variable susceptibility to paired drugs previously implicated in collateral sensitivity (for selection drugs cefuroxime, chloramphenicol, gentamicin, streptomycin and trimethoprim, the paired antibiotics were gentamicin^6^, polymyxin B^6^, cefuroxime^6^, tetracycline^5^ and nitrofurantoin^5^). Unlike resistance to selection drugs, where there was an average increase in resistance relative to the ancestral strain (mean ±s.d. of log2 transformed relative IC_90_: 1.78±1.27, Fig. 2), the average fold change in IC_90_ to paired drugs was lower (0.12±0.97, Fig. 3) and encompassed both positive (cross resistance) and negative (collateral sensitivity) changes in resistance. Average collateral resistance varied depending on which gene was mutated for mutants selected against gentamicin then tested against cefuroxime (Fig. 3c, F_5,126_ = 11.2, p<0.0001) and mutants selected against trimethoprim and tested against nitrofurantoin (Fig. 3c, χ^2^_4_ = 10.0, p<0.05), but not for the other three drug pairs (p>0.05 in all cases). We found several genes that on average induced collateral sensitivity to paired drugs, such as *ubi* mutants which induced collateral sensitivity to cefuroxime (effect of *ubi* mutation on log_2_ transformed relative IC_90_: β = −1.27, t_137_=5.68, p<0.0001) and *atpG* mutants which led to collateral sensitivity to tetracycline (β = −1.27, t_60_=2.31, p<0.05). Mutations in *ubi* genes and in *atpG* affect ubiquinone synthesis and the ATP synthase respectively disrupting the proton motive force (PMF), leading to a reduced membrane potential and hence reduced influx of aminoglycosides^26^. Despite the benefit of aminoglycoside resistance, PMF-driven efflux pumps such as acrAB are less active in mutants with disrupted PMF^5^, increasing susceptibility to other drugs.

**Figure 3:**
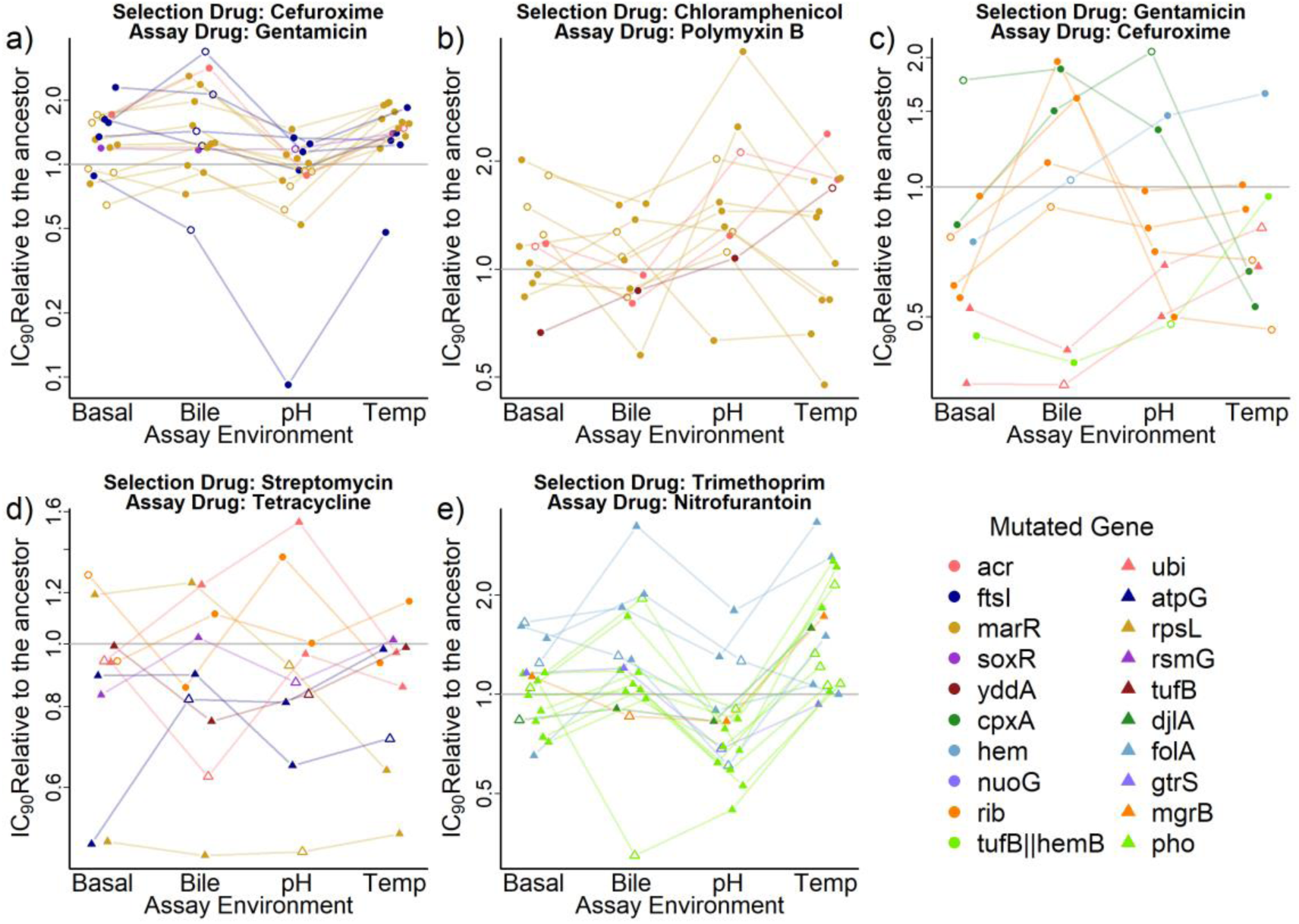
Collateral changes in resistance to a paired drug (*assay drug*) for resistant mutants selected with each *selection drug* (panels a-e), tested across different abiotic conditions (*x*-axis, *assay environments*). Each set of four connected points shows a single resistant mutant, coloured according to which gene was mutated (see legend). Resistance is shown relative to the IC_90_ of the ancestral strain in the same assay environment (scores for the ancestor are shown in Fig. S1). Hollow points indicate sympatric combinations (selection environment = assay environment). Each point is the mean of 2-4 independent replicates (mean *n* = 3.83). The *y* axis is log transformed and the scale varies between panels.

Because selection environment influenced which genes were mutated (Fig. 1), and in turn mutated gene influenced collateral resistance phenotypes to paired drugs (Fig. 3), we tested whether this translated to variation of mean resistance to paired drugs depending on selection environment. We found such an effect for mutants selected against streptomycin and assayed against tetracycline (χ^2^_3_ = 8.58, p<0.05). Here only mutants selected in the bile (relative IC_90_ of bile selected mutants β=- 0.34, t_36_=2.25,p<0.05) and low pH (β=-0.35, t_36_=2.25,p<0.01) environments showed significant collateral sensitivity. We also found a significant effect of selection environment on resistance to nitrofurantoin for mutants selected against trimethoprim (effect of selection environment: χ^2^_3_ = 12.6, p<0.01). For these trimethoprim-selected mutants, *folA* mutations were more common in basal and pH selection environments (Fig. 1), and had increased nitrofurantoin resistance (effect of *folA* mutation on log_2_ transformed relative IC_90_: β = 0.40, t_111_=2.81, p<0.01). By contrast *phoPQ* mutants, which did not have significantly altered nitrofurantoin resistance (β=-0.07, t_146_=0.64,p>0.5), were more common in the bile and high temperature selection environments. This shows that for some of the antibiotics and environments we tested, variation of the abiotic conditions during resistance evolution to one antibiotic resulted in variation of average collateral resistance to other antibiotics.

Expression of average collateral sensitivity / cross resistance to paired drugs also varied depending on the abiotic conditions during exposure to the paired drug. For mutants selected against cefuroxime, chloramphenicol and trimethoprim, there was a significant effect of assay environment on average resistance to the paired drug, relative to the ancestor in the same conditions (effect of assay environment for selection/paired drug: cefuroxime/gentamicin, χ^2^_3_ = 44.6, p<0.0001; chloramphenicol/polymyxin B, χ^2^_3_ = 10.7, p<0.05; trimethoprim/nitrofurantoin, χ^2^_3_ = 64.7, p<0.0001). Note this variation of susceptibility relative to the ancestral strain was not explained by variable susceptibility of the ancestral strain across assay environments (Fig. S1). Qualitatively similar results emerge (Fig. S1) if we use the absolute IC_90_ value (not relative to the ancestor).

Changing the local abiotic conditions did not affect all mutants the same way: the mutated gene and assay environment interacted to determine resistance to paired drugs for three of the drug pairs (Genotype by assay environment interaction for selection/paired drug: cefuroxime/gentamicin, χ^2^_9_ = 19.3, p<0.05; chloramphenicol/polymyxin B, χ^2^_6_ = 14.4, p<0.05; gentamicin/cefuroxime, χ^2^_9_ = 34.1, p<0.001). In some cases this variation led to a switch between cross resistance and collateral sensitivity, such as *cpxA* mutants which were resistant to cefuroxime in the bile and pH environments but were susceptible to cefuroxime at high temperature (Fig. 3c). *cpxA* is part of a two-component regulator which responds to misfolded proteins in the periplasm, activating the cpx response, which has been shown to confer resistance to aminoglycosides^32^. Mutations in the periplasmic domain of *cpxA* (as in our mutants) have the cpx pathway locked into an activated state^33^, with cpx phenotypes being more pronounced at high temperature ^34,35^. Other work has shown that, due to its influence on cell wall homeostasis, the cpx response can influence resistance to β-lactams like cefuroxime, but that it must be at an intermediate level for maximal resistance^36^. At temperatures of 37°C our mutants likely upregulate the cpx response into the optimum zone, leading to β-lactam resistance. However at 42°C the mutant’s cpx response is likely further upregulated^34,35^, meaning that peptidoglycan homeostasis is no longer maintained, resulting in β-lactam sensitivity^36^.

In summary, mutants selected for resistance to one antibiotic often had altered resistance to other antibiotics (as expected, having chosen these antibiotics based on past evidence of such effects). However, these collateral effects varied among different pathways to resistance (mutated genes), which translated to variation in average collateral effects depending on the abiotic conditions during selection for resistance to the first antibiotic (selection environment). Finally some collateral effects were only pronounced in specific environmental conditions.

### Antibiotic-free growth depends on resistance mechanism and local abiotic conditions

Antibiotic resistance is frequently associated with a cost in terms of impaired growth in the absence of antibiotics, and variation of this cost is a key driver of the long- term persistence of resistance^37^. In our mutant-selection experiment bacteria were grown in the absence of antibiotics prior to plating. Therefore, variable costs of resistance across selection environments could potentially have influenced which mutations we detected in which sets of conditions. We investigated this by quantifying the growth capacity of each mutant in the absence of antibiotics, using the same dataset used to calculate the IC_90_ values (see methods). Relative to the ancestral strain in the absence of antibiotics (mean final optical density ± standard deviation: 0.90 ± 0.16), most types of resistant mutants showed evidence of average growth costs, although this varied depending on the selection drug (gentamicin 0.62±0.34; streptomycin 0.49±0.29; cefuroxime 0.89±0.21; chloramphenicol 0.82±0.15; trimethoprim 0.80±0.21).

For all selection drugs, mean antibiotic-free growth varied depending on which gene was mutated (effect of mutated gene on antibiotic-free growth: cefuroxime-selected mutants, F_4,17_= 128, p<0.0001; chloramphenicol, F_3,70_ = 601, p<0.0001; gentamicin, F_6,12_ = 30.6, p<0.0001; streptomycin, F _6,10_ = 14.9, p<0.0001; trimethoprim, F _5,19_ = 256, p<0.0001). For example, streptomycin-resistant mutants with mutations in *atpG* grew relatively poorly (Fig 4d). For trimethoprim-resistant mutants, this variation translated to differences in average antibiotic-free growth among sets of mutants from different selection environments (effect of selection environment: χ^2^_3_ = 17.3, p<0.001; Fig. 4e). With this antibiotic, the mutants selected at high temperature and included in our phenotyping assay (in 5/5 cases, *phoPQ* mutants, Table S1) grew relatively poorly across all antibiotic-free environments compared with other mutants (Fig. 4e). However, when assayed at high temperature the *phoPQ* mutants were on average more resistant (Fig. 2e and Fig. S3e) than other types of mutants. This suggests that *phoPQ* mutants were less frequently isolated from selection environments at 37°C because the cost of the resistance mutation was not counterbalanced by more effective resistance in these selection environments. This is also consistent with the local adaptation we saw above for these mutants in our analysis of their growth at selection concentration (GASC).

**Figure 4:**
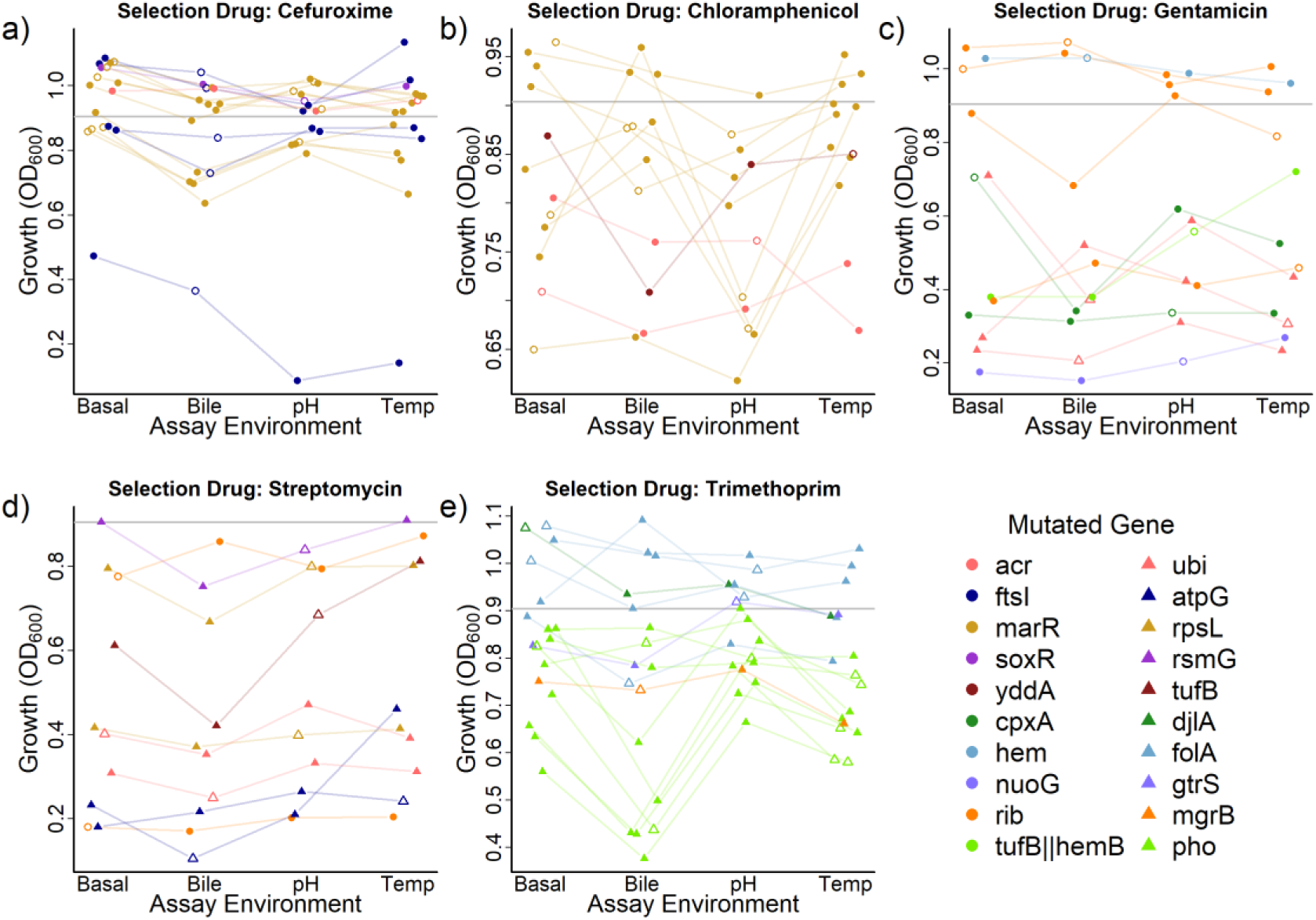
Antibiotic-free growth of resistant mutants selected with each *selection drug* (panels a-e), measured in four different *assay environments*. Each set of four connected points shows a single resistant mutant, coloured according the gene or gene family that was mutated. Growth was measured in the absence of antibiotics as optical density (OD600) after 20h. Hollow points indicate sympatric combinations (selection environment = assay environment). The grey line shows the antibiotic-free growth of the ancestral strain. Points are means of 2-4 independent replicates (mean *n* = 3.89). The *y* axis varies between panels.

For all selection drugs, variation of antibiotic-free growth among different genotypes depended on the local abiotic conditions (genotype by assay environment interaction: cefuroxime: χ^2^_9_ = 24.0, p>0.01; chloramphenicol: χ^2^_6_ = 26.6, p<0.001; gentamicin: χ^2^_15_ = 43.6, p<0.001; streptomycin: χ^2^_15_ = 36.8, p<0.01 and trimethoprim χ^2^_12_ = 63.1, p<0.0001). Despite this, we did not find any evidence of local adaptation in terms of antibiotic-free growth (p>0.05 for all selection drugs).

### Conclusions

Our findings have important implications for research that aims to exploit collateral sensitivity in novel treatment approaches. For example, among our streptomycin-resistant mutants, only those selected in the bile and pH environments showed significant collateral sensitivity. Thus, streptomycin treatment of enteric *E. coli* (exposed to low pH and bile) could potentially select for mutants that are collaterally sensitive to tetracycline, but this may be less likely in other environments. Studies of collateral sensitivity will therefore be most relevant when they account for environmental variation, for example by focusing on resistant mutants that arise under conditions similar to those during infection, or testing explicitly for variation across abiotic conditions (as above). This will ensure collateral effects of mutations specific to the infection environment are not overlooked, and that collaterally sensitive mutants specific to the lab environment are not given undue attention. Note our study included only chromosomal mutants derived from a single strain, albeit of an important commensal and opportunistically pathogenic species. Nevertheless, some of the resistance mechanisms we identified are known to be important in natural and clinical populations, such as mutations in genes for efflux pumps (*acr*)^38^, global regulators (marR, *phoPQ*)^39–41^ and specific antibiotic targets (*ftsI* and *rpsL*)^42–44^, suggesting our findings are relevant beyond laboratory studies. A key question for future work is whether collateral effects of resistance encoded on plasmids^45,46^, which is common in clinics, show similar sensitivity to abiotic conditions as we saw here. We speculate this is likely, because plasmids can carry accessory genes with fitness effects that are strongly affected by the abiotic environment ^47,48^.

To ensure that clinically relevant mutants are assayed for collateral sensitivity, several past studies have looked at clinical isolates directly^10,49,50^. However, these studies each measured collateral sensitivity in a single lab environment. Our results suggest this can potentially lead to overlooking collateral sensitivity interactions that occur in the infectious environment. For example, if we had only tested *cpxA* mutants at 37C°C, as has been done previously^51,52^, we would have only observed that they mediate cross resistance between gentamicin and cefuroxime. This would miss the collateral sensitivity of these mutants at higher temperature, a potentially interesting target for collateral sensitivity in *E. coli* infecting hosts with higher body temperatures, e.g. poultry^24^. The changing expression of collateral effects with abiotic conditions therefore represents an important variable which should be considered if we are to design robust treatment regimens around collateral sensitivity. Of course, our experiments were still restricted to simplified lab conditions. Our aim here was to demonstrate that even relatively minor (but still relevant to nature) manipulations of the abiotic environment can modify collateral sensitivity interactions. Our finding that such effects are strong suggests other differences between complex within-host or natural environments and in vitro screening conditions are likely to modify expression of collateral resistance even further.

In summary, we found local abiotic conditions modified collateral sensitivity interactions by two principal mechanisms. First, antibiotic treatment in different abiotic environments selects for different pathways to resistance (Fig. 1), and these are sometimes associated with variable collateral effects. For the three selection drugs where we found selection environment influenced which mutations occurred (cefuroxime, streptomycin and trimethoprim), the mutants selected in different environments had different resistance phenotypes, both to the selection drug and collateral effects to other drugs. Second, for individual mutants, expression of collateral sensitivity or cross resistance can depend strongly on local abiotic conditions. For example, we found some gentamicin-resistant mutants were cross resistant or collaterally sensitive to cefuroxime depending on the assay environment. This is consistent with a more general trend that the phenotypic effects of antibiotic resistance mechanisms are highly sensitive to genotype by environment interactions ^16–18,53^. Critically, this suggests for some antibiotic combinations, the effectiveness of the second drug against bacteria that have evolved resistance to the first drug, and consequently selection on resistance to both drugs, will depend on the abiotic environment. While collateral sensitivity still holds great promise as a way to prolong the effectiveness of available treatments, we suggest doing so will be most effective if we account for local abiotic conditions.

## Methods

### Organisms and growth conditions

We used *Escherichia coli* MG1655 as the ancestral organism, grown at 37°C in static 100μl cultures in 96-well microplates unless otherwise stated. The media used was based around lysogeny broth (LB, Sigma Aldrich) with additions to the media to create variation in environmental conditions for basal, pH, and bile media. Basal media (the base condition) is LB buffered at pH 7.0 with 0.1M sodium hydrogen phosphate (Na_2_HPO_4_ and NaH_2_PO_4_). pH media (acidic pH), is LB buffered at pH 6.5 with 0.1M sodium hydrogen phosphate. Bile media is LB with the addition of 0.5 g/L of bile salts and buffered at pH 7.0 with 0.1M sodium hydrogen phosphate. The temperature treatment uses the basal media but is incubated at 42°C. When these conditions were on solid media (i.e. to select for resistant colonies) we instead used LB agar (Sigma Aldrich), but other temperature and media additions were the same. For overnight culture prior to assay we incubated at 37°C in diluted LB (1:2 LB:water).

### Mutant isolation

We screened for mutants resistant to each selection drug in each selection environment by first culturing the ancestral *E. coli* strain in each of the 4 environments for 20 hours in the absence of antibiotics. After 20 hours growth, each entire culture was transferred to a well of a 24-well plate containing 1ml of agar corresponding to the same environment as the prior liquid culture plus one of five antibiotics at selection concentration (cefuroxime [selection concentration = 6 μgml^- 1^], chloramphenicol [6 μgml^-1^], gentamicin [24 μgml^-1^], streptomycin[72 μgml^-1^] and trimethoprim [0.5 μgml^-1^]). These selection concentrations were approximately equal to minimal inhibitory concentrations for the ancestor. This meant the ancestral strain was effectively killed, but mutants with even moderately increased resistance could grow. A higher selection concentration was used for the aminoglycosides (gentamicin, 48 μgml^-1^; streptomycin, 144 μgml^-1^) in the pH media because of the reduced efficacy against the ancestral strain in this environment ^27^. The agar with bacterial culture was incubated for 48 hours at 37 or 42°C. This protocol was used to test 92 independent *E. coli* populations against each combination of 4 selection environments and 5 selection drugs (1840 populations total). After incubation, each agar well was checked for the appearance of resistant colonies (the number of populations that produced colonies is shown in figure S4). Up to 6 colonies for each drug by environment combination were picked from 6 randomly selected, independent agar wells (populations). These colonies were grown in LB without antibiotics for 3 hours before glycerol was added to 25% of the final volume and mutants were frozen at −80°C. Mutant isolation in the basal, pH and temperature environments was done in the same temporal block, and the bile environment in a separate block. For a minority of mutants, we were unable to consistently revive the frozen stock for library preparation for sequencing and/or for phenotyping. These were excluded from the relevant assays and analysis (see Table S1 for details).

### Genome sequencing and bioinformatics

Genomic DNA from 110 mutants plus the ancestral strain was extracted with the Genomic-tip 20/G Kit (Cat. No. 10223, Qiagen) according to the manufacturer’s instructions. Libraries were produced using the Illumina Nextera XT kit. Sequencing was performed on the Illumina HiSeq 4000 platform with 150 bp paired end reads at the Functional Genomic Center, Zürich, Switzerland (Fig. 1). Reads were trimmed using trimomatic^54^ and then analysed using the breseq pipeline^55,56^. Each mutant should be clonal with respect to the mutations contributing to resistance, so we only considered mutations that were at frequency of greater than 75%. For mutants that had putative resistance mutations at a frequency 20%-75%, we replica plated 20 colonies with and without the selective conditions (selection environment + selection drug). Where the frequency of resistant colonies was consistent with the frequency of the putative resistance mutation, we picked one resistant colony (from the non-selective plate), re-cultured it and used it in phenotyping, otherwise the mutant was not used for further analysis. Finally, several strains did not have high-frequency mutations which could be attributed to resistance, so were not used for further analysis. Once we had completed these filtering steps, we gained full genotypic information for 95 strains.

### Measuring resistance to selection drugs and other drugs

So as to make the phenotyping manageable, we selected 85 mutants (and the ancestor) to phenotype, excluding mutants where very similar genotypes were already represented (Table S1). Resistance of all mutants and the ancestral strain was quantified using broth dilution. For each antibiotic, we assayed each combination of strain, assay environment and antibiotic concentration in four replicates, each in a separate temporal block. In each block of assays, we used a frozen masterplate containing all strains organized in one of three randomized layouts (blocks 1, 2, 3 and 4 used layouts 1, 2, 3 and 1) to inoculate a single pre- culture plate (1:2 LB:water). We then incubated the pre-culture plate for 3 hours before using it to inoculate all the overnight plates (for every assay culture we grew a separate overnight culture). Overnight cultures were then used to inoculate the assay plates using a pin replicator. Each mutant was tested against the relevant selection- and paired- drugs, and the ancestor was tested against all drugs, each at 8 concentrations (including zero) in each assay environment. After culturing assay plates for 20 hours, we agitated the plates to resuspend bacteria, then measured the biomass of bacteria by optical density at 600nm (OD_600_) using a spectrophotometer (Infinite®200 PRO, Tecan Trading AG, Switzerland). Due to the time taken to read 64 plates, incubation and plate reading was staggered and the order was randomised. Some mutants failed to regrow during overnight incubation, resulting in false inoculation of some assay wells. To filter out these false negatives, we excluded OD_600_ scores from assay plates that were <0.075, but only if the OD_600_ in the overnight well (prior to inoculation of the assay well) was also <0.075.

### Calculation of summary phenotypes from dose response data

For each mutant strain we calculated 4 phenotypes 1) 90% inhibitory concentration (IC_90_) to the selection drug, 2) growth at selection concentration (GASC) for the selection drug, 3) IC_90_ to the paired drug and 4) growth in absence of antibiotics. These phenotypes were calculated from the dose response relationship data for the selection and paired drugs (for each replicate separately). We fitted a Hill function using non-linear least squares in R^57^:

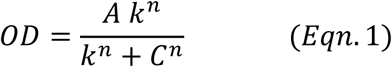

Where OD is the measured optical density and C is the drug concentration. *A* is then the asymptote, *k* is the inflection point of the curve and *n* is the Hill parameter controlling curve steepness. Thus, growth in the absence of antibiotics is equal to *A*, and for each combination of mutant and assay environment is the mean of the A parameters for all dose-response curves. The IC_90_ for the selection and paired drugs can be calculated using the following formula taking the parameters from the relevant fitted curve.

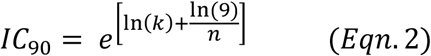

Finally, the growth at selection concentration (GASC) is the value of the Hill function when the antibiotic concentration equals the selection concentration.

For some strain : drug : environment combinations we could not robustly fit a Hill function to some or all replicates, for example if there was very little growth inhibition or if the dose response was strongly stepwise. In these cases, where possible, we calculated phenotypes independently from the fitting of dose response curves (as is often done in other studies^6^). We estimated growth in the absence of antibiotics from the OD in the absence of antibiotics. We took the IC_90_ as the lowest tested concentration where growth was below 10% of growth in the absence of antibiotics. Finally, we took growth at selection concentration as the OD score at the selection concentration of the antibiotic, or the predicted score at the selection concentration assuming a linear relationship between growth and drug concentration between the two measured concentrations on either side of the selection concentration. We used these fit-independent methods for a minority of cases (9.7% for antibiotic free growth, 5.3% for selection drug IC_90_, 7.6% for pair drug IC_90_, and 8.6% for GASC). In all cases there was a strong correlation between the fit-dependent and fit-independent measures (Fig S5).

### Statistics

We treated the mutants selected for resistance against different antibiotics as five independent data sets, due to the difficulty in comparing resistance across multiple antibiotics. In each dataset, we took the four phenotypes of interest (see above) as response variables in separate models. In each model, the replicate measures for each phenotype came from independent dose response curves (fitted to data collected in different blocks). Mutants with insufficiently replicated data (<2 replicates in any of the four assay environments) for a given phenotype were excluded from the analysis for that phenotype (Table S1). We transformed IC_90_ values by taking log_2_(IC_90_) relative to the mean of the ancestral strain measured in the same environment. This controlled for any effects of assay environment on antibiotic-inhibition of the ancestral strain (Fig. S1) and normalised the data. GASC was square-root transformed to fit the assumption of normality, but was not relative to the ancestor as the GASC was not significantly different from zero in the ancestor, as expected. Growth in the absence of antibiotics already fit the assumptions of normality was not analysed relative to the ancestor because the ancestral growth did not vary significantly between assay environments (χ^2^_3_ = 5.95, p>0.05).

Then for each of the 4 phenotypes across the 5 datasets we fitted two mixed effects models (Table S2). The fixed effects of these two models were:

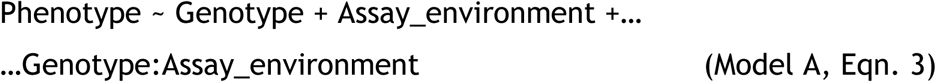

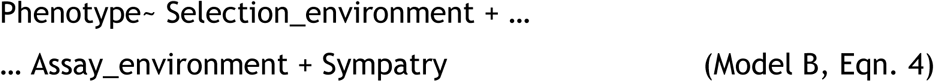

Where genotype is based on the gene or operon mutated (so that mutants with different mutations in the same gene/operon have the same genotype), and sympatry is a binary vector indicating whether selection environment is the same as assay environment. Model A was used to test whether genotypes varied in their average phenotypes (effect of genotype) and whether that variation depended on the local abiotic conditions (genotype by assay environment interaction). To test the effect of selection environment, we used a separate model (B) because genotype and selection environment were often confounded. Model B was used to test whether each phenotype varied on average among sets of mutants isolated from different selection environments (effect of selection environment), and for evidence of local adaptation in terms of higher average phenotypic scores in sympatric compared to allopatric combinations (main effect of sympatry, where sympatry means selection environment = assay environment, and allopatry means selection environment ≠ assay environment)^31^.

In both models we included a nested random effect of strain (individual mutant ID) on intercept (+(1|Strain) in the lmer function) to account for variation between strains^58^. We also included a random effect of block nested within strain on the intercept (+(1|Strain:Block) in the lmer function), to account for variation between measures of the same strain in different blocks. To prevent overfitting, the variance explained by these random effects was tested using a likelihood ratio test (on the maximal model) and non-significant terms were dropped, potentially reducing to fixed effects model if both random effects were dropped.

Significance of terms in models is reported from minimal models by comparing models with or without the term of interest using a likelihood ratio test (χ^2^ statistic), except when the main effect of genotype is involved in a significant higher order interaction (genotype: assay_environment). In these cases, the main effect cannot be dropped, so significance is instead reported with an F test (on a type III ANOVA), using the approximate degrees of freedom calculated using lmerTest^59^. Although assay environment is included in both models, we report the significance of the main effect of assay environment from model B, where it does not have higher order interactions and can always be tested using a likelihood ratio test.

To test for significant differences between mutations acquired by mutants selected in the four selection environments, we used a permutational ANOVA on the data for which genes were affected by the mutations (Table 2, full genotype). This was performed using the adonis function in the vegan package^60^.

## Acknowledgements

RCA was funded by the a postdoctoral fellowship from ETH Zurich and Marie Curie Actions for People (FEL-28 16-1). ARH was funded by SNSF Project 31003A_165803.

