## Supplemental Figures S1-S5 and tables S1 & S2 for "Collateral sensitivity interactions between antibiotics depend on local abiotic conditions"

### Supplementary information for: Collateral sensitivity interactions between antibiotics depend on local abiotic conditions

#### Supplementary Figures

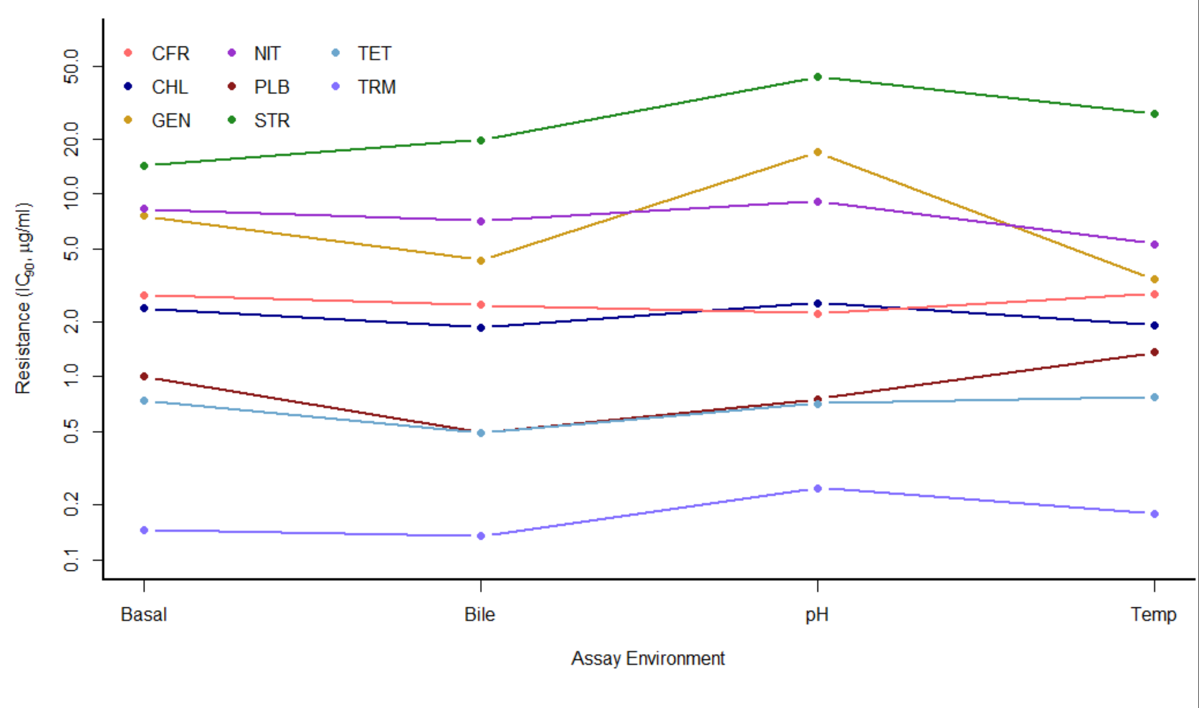


**Fig. S1: $IC_{90}$ for the ancestral *E. coli* strain to each of the antibiotics used in this study, assayed in each of the different environments.** Points are the mean of 2-4 replicates and the y axis is log transformed. Assay environment had a predictive effect on the level of resistance to gentamicin ($\chi^2_3$ = 29.5, p<0.0001), polymyxin ($\chi^2_3$ = 10.8, p<0.05) and streptomycin ($\chi^2_3$ = 11.8, p<0.01). All $IC_{90}$ comparisons reported in the text are relative to the ancestral $IC_{90}$ in the same abiotic environment. This could in theory influence the main effect of assay environment on collateral effects as reported in the main text (seen for the following selection/paired drug combinations of cefuroxime/gentamicin, chloramphenicol/polymyxin B and trimethoprim/nitrofurantoin). However, we see the same main effect of assay environment in two of these three combinations when we use absolute $IC_{90}$ instead of IC90 relative to the ancestor (cefuroxime/gentamicin, $\chi^2_3$ = 141, p<0.0001 ; chloramphenicol/polymyxin B $\chi^2_3$ = 69.4, p<0.0001). We also see a significant effect for streptomycin selected mutants (streptomycin / tetracycline $\chi^2_3$ = 20.1, p<0.001), which was not significant using IC90 relative to the ancestor (see main text).

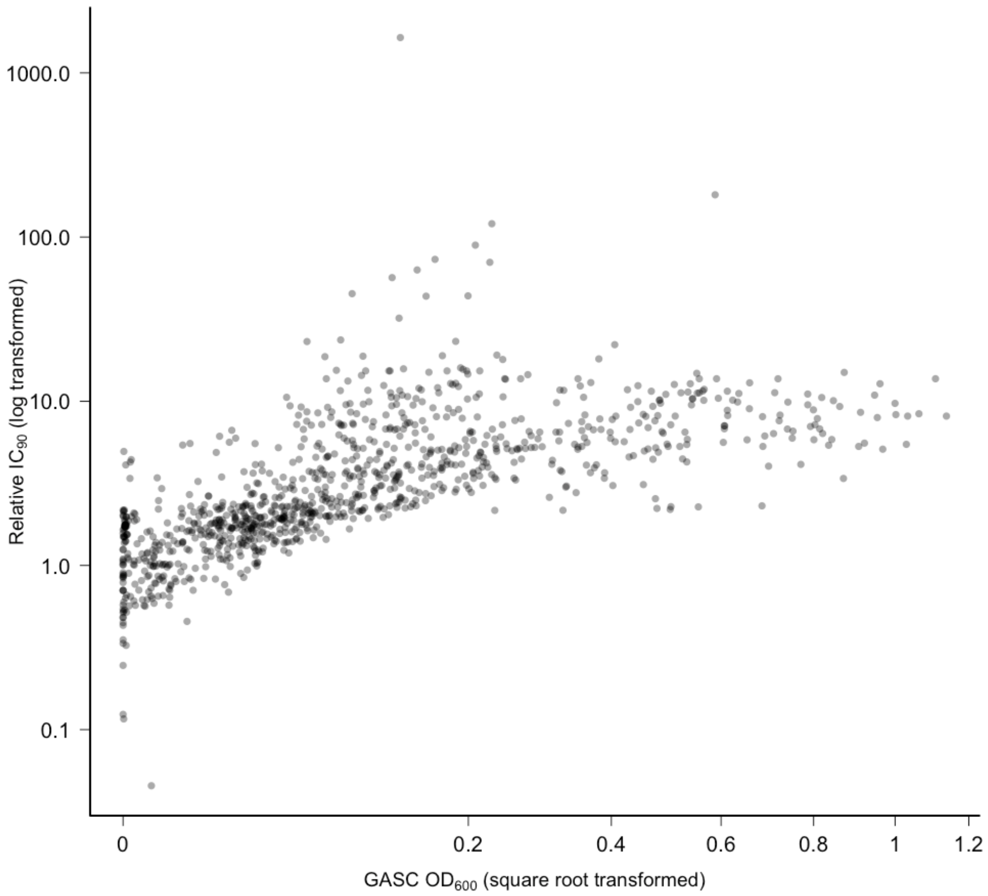


**Figure S2: Correlation between growth at selection concentration (GASC) and $IC_{90}$ for the ancestor and 66 mutants, each measured against their relevant selection drug in each of the four assay environments**. Each point is from an independent dose response assay, with $IC_{90}$ and GASC calculated from the same data. The different measures are transformed according to the way that the variables are input into the relevant models. These measures are highly correlated ($\tau$=0.61, p<0.0001).

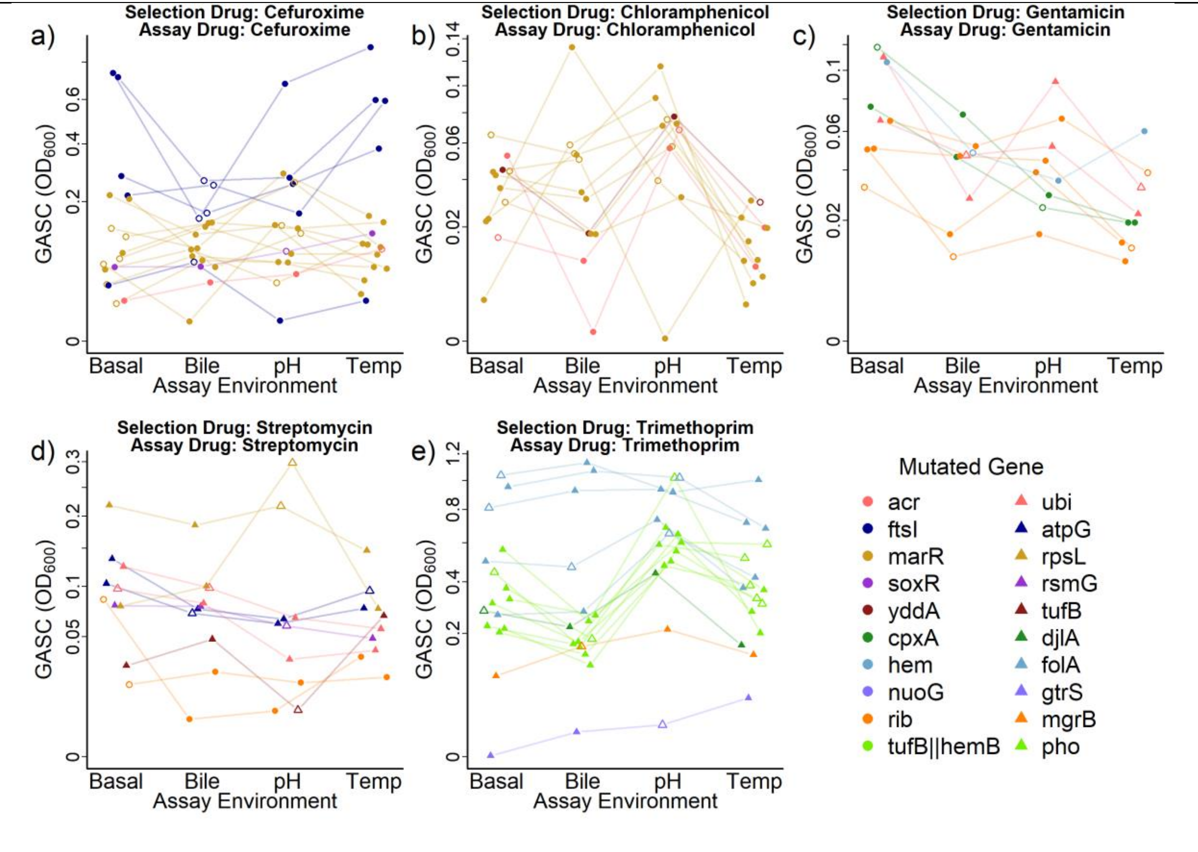


**Fig. S3: Population growth for resistant mutants selected with each selection drug (panels a-e) when grown with that antibiotic at selection concentration (GASC), measured in four different assay environments.** Each set of four connected points shows a single resistant mutant, coloured according the gene or gene family that was mutated. Growth was measured in the presence of the selection drug at the selection concentration (see methods) using optical density (OD600) after 20h. Hollow points indicate sympatric combinations (selection environment = assay environment). Points are means of 2-4 independent replicates (mean n = 3.86). The y axis is square-root transformed and varies between panels. GASC varied depending on mutated gene (main effect of genotype: cefuroxime, $F_{4,20} = 21.7$, $p<0.0001$; chloramphenicol, $F_{3,79} =24.7$, $p<0.0001$; gentamicin, $\chi^2_3 = 9.07$, $p<0.05$; streptomycin, $F_{11,134} = 181$, $p<0.0001$ and trimethoprim, $F_{5,19}=83.6$, $p<0.0001$), and the interaction between mutated gene and assay environment (genotype by assay environment interaction effect on GASC: cefuroxime, $\chi^2_9 = 21.0$, $p<0.05$; chloramphenicol, $\chi^2_6 = 13.1$, $p<0.05$; streptomycin, $\chi^2_{15} = 26.3$, $p<0.05$; and trimethoprim, $\chi^2_{12} = 47.6$, $p<0.0001$).

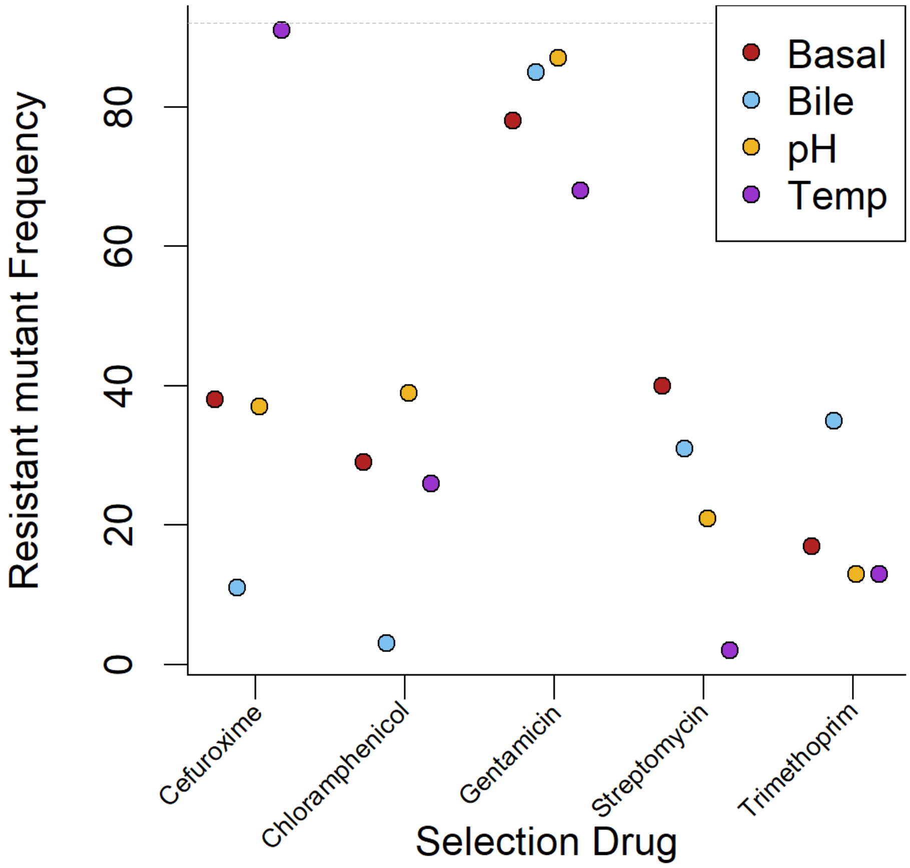


**Figure S4: Number of independent wells (out of 92) which had resistant colonies after plating *E. coli* with different drugs in different environments.** The number of populations in which resistance was seen (one or more resistant colonies) varied with the combination of antibiotic and selection environment (binomial glm of number of independent wells with resistance: Antibiotic : environment interaction, $\chi^2_{12} = 160$ $p<0.0001$).

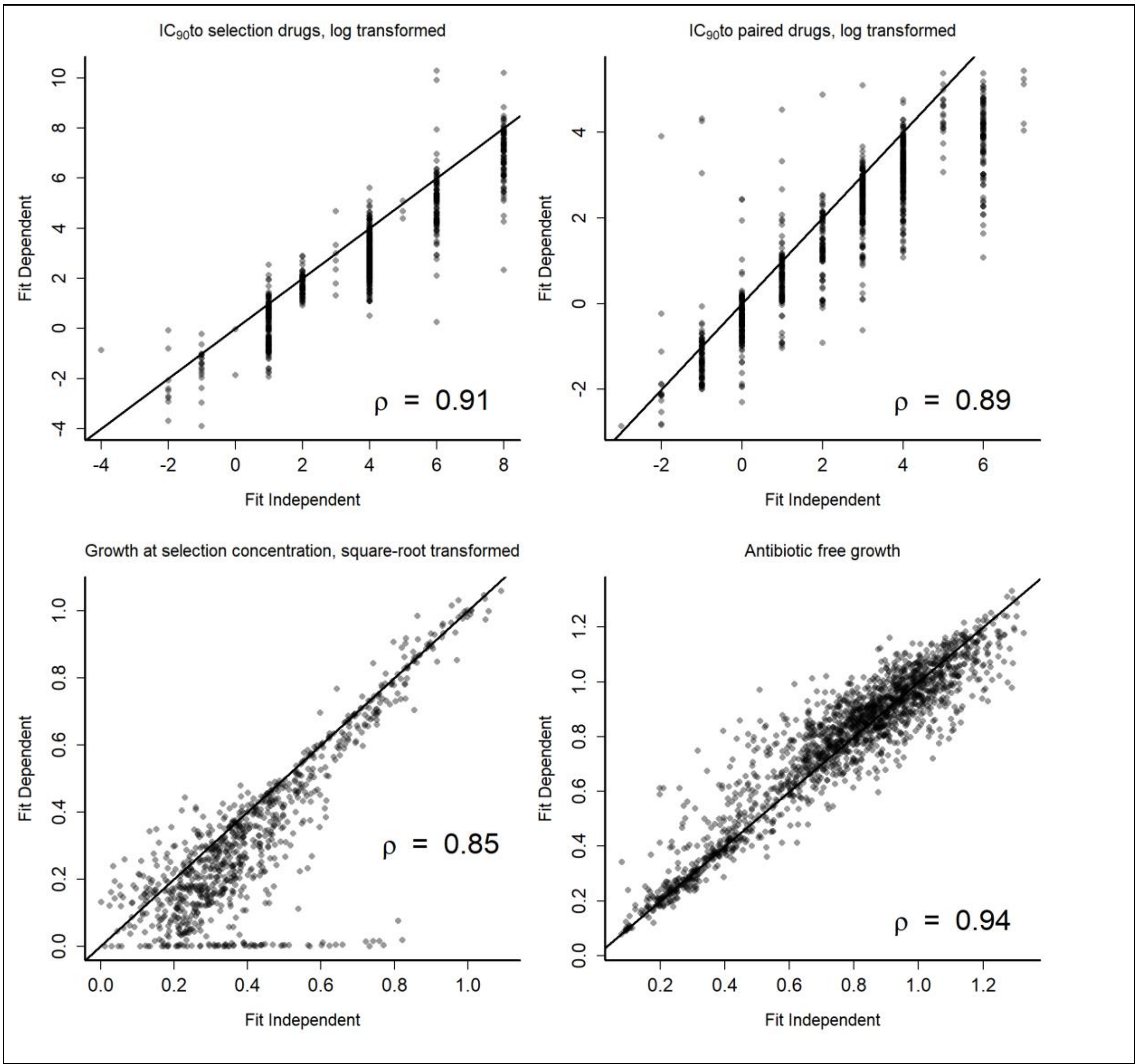


**Fig S5: Correlation between the 4 phenotypes measured when calculated using a method that requires a fitted Hill function (fit dependent) and a method that does not require a fitted Hill function (fit independent).** Points represent values calculated from individual replicate dose response measurements. Only points where both fit dependent and fit independent methods were calculated are plotted. The correlation measured by Pearson's correlation coefficient is given at the bottom-right of each panel.

#### Supplementary Tables

| Selection Drug | Selection Environment | Replicate Mutant | Genotype | | Included in phenotype assays | Included in model | | | | Notes |
|---|---|---|---|---|---|---|---|---|---|---|
| | | | Full | Family | | Selection drug $IC_{90}$ | GASC | Paired drug $IC_{90}$ | Antibiotic Free growth | |
| Cefuroxime | Basal | 1 | marR | mar | Yes | Yes | Yes | Yes | Yes | |
| Cefuroxime | Basal | 2 | marR | mar | Yes | Yes | Yes | Yes | Yes | |
| Cefuroxime | Basal | 3 | marR | mar | Yes | Yes | Yes | Yes | Yes | |
| Cefuroxime | Basal | 4 | marR | mar | Yes | Yes | Yes | Yes | Yes | |
| Cefuroxime | Basal | 5 | marR \|\| yegH/asmA | mar | Yes | Yes | Yes | Yes | Yes | |
| Cefuroxime | Basal | 6 | marC/marR | mar | Yes | Yes | Yes | Yes | Yes | |
| Cefuroxime | Bile | 1 | marR | mar | Yes | No | No | No | No | |
| Cefuroxime | Bile | 2 | ftsI | ftsI | Yes | Yes | Yes | Yes | Yes | |
| Cefuroxime | Bile | 3 | ftsI | ftsI | Yes | Yes | Yes | Yes | Yes | |
| Cefuroxime | Bile | 4 | ftsI | ftsI | Yes | Yes | Yes | Yes | Yes | |
| Cefuroxime | Bile | 5 | ftsI \|\| yraR | ftsI | Yes | Yes | Yes | Yes | Yes | |
| Cefuroxime | Bile | 6 | ftsI | ftsI | Yes | Yes | Yes | Yes | Yes | |
| Cefuroxime | pH | 1 | marR | mar | Yes | Yes | Yes | Yes | Yes | |
| Cefuroxime | pH | 2 | marR | mar | Yes | Yes | Yes | Yes | Yes | |
| Cefuroxime | pH | 3 | soxR | soxR | Yes | Yes | Yes | Yes | Yes | |
| Cefuroxime | pH | 4 | | | No | No | No | No | No | No mutation identified |
| Cefuroxime | pH | 5 | marR | mar | Yes | Yes | Yes | Yes | Yes | Enriched |
| Cefuroxime | pH | 6 | pykF \|\| marR | mar | Yes | Yes | Yes | Yes | Yes | |
| Cefuroxime | Temperature | 1 | | | No | No | No | No | No | No mutation identified |
| Cefuroxime | Temperature | 2 | | | No | No | No | No | No | No mutation identified |

| | | | | | | | | | | |
|---|---|---|---|---|---|---|---|---|---|---|
| Cefuroxime | Temperature | 3 | | | No | No | No | No | No | No mutation identified |
| Cefuroxime | Temperature | 4 | | | No | No | No | No | No | No mutation identified |
| Cefuroxime | Temperature | 5 | acrR | acr | Yes | Yes | Yes | Yes | Yes | |
| Cefuroxime | Temperature | 6 | | | No | No | No | No | No | No mutation identified |
| Chloramphenicol | Basal | 1 | marR | mar | No | No | No | No | No | Duplicated with CHL3Bile |
| Chloramphenicol | Basal | 2 | | | No | No | No | No | No | No mutation identified |
| Chloramphenicol | Basal | 3 | marR | mar | Yes | Yes | Yes | Yes | Yes | Enriched |
| Chloramphenicol | Basal | 4 | acrR | acr | Yes | Yes | Yes | Yes | Yes | Enriched |
| Chloramphenicol | Basal | 5 | marR | mar | Yes | Yes | Yes | Yes | Yes | Enriched |
| Chloramphenicol | Basal | 6 | marC/marR | mar | Yes | Yes | Yes | Yes | Yes | Enriched |
| Chloramphenicol | Bile | 1 | marR | mar | Yes | Yes | Yes | Yes | Yes | |
| Chloramphenicol | Bile | 2 | marR | mar | Yes | Yes | Yes | Yes | Yes | |
| Chloramphenicol | Bile | 3 | marR | mar | Yes | Yes | Yes | Yes | Yes | |
| Chloramphenicol | pH | 1 | marR \|\| wzxB | mar | Yes | Yes | Yes | Yes | Yes | |
| Chloramphenicol | pH | 2 | | | No | No | No | No | No | No mutation identified |
| Chloramphenicol | pH | 3 | marR | mar | Yes | Yes | Yes | Yes | Yes | |
| Chloramphenicol | pH | 4 | marR | mar | Yes | Yes | Yes | Yes | Yes | Enriched |
| Chloramphenicol | pH | 5 | acrA/acrR | acr | Yes | Yes | Yes | Yes | Yes | |
| Chloramphenicol | pH | 6 | | | No | No | No | No | No | No mutation identified |
| Chloramphenicol | Temperature | 1 | | | No | No | No | No | No | No mutation identified |
| Chloramphenicol | Temperature | 2 | | | No | No | No | No | No | No mutation identified |

| | | | | | | | | | | |
|---|---|---|---|---|---|---|---|---|---|---|
| Chloramphenicol | Temperature | 3 | | | No | No | No | No | No | No mutation identified |
| Chloramphenicol | Temperature | 4 | yddA | yddA | Yes | Yes | Yes | Yes | Yes | |
| Chloramphenicol | Temperature | 5 | | | No | No | No | No | No | No mutation identified |
| Chloramphenicol | Temperature | 6 | | | No | No | No | No | No | No mutation identified |
| Gentamicin | Basal | 1 | ribB/yqiC | rib | Yes | Yes | Yes | Yes | Yes | |
| Gentamicin | Basal | 2 | cpxA | cpxA | Yes | No | No | No | No | |
| Gentamicin | Basal | 3 | cpxA | cpxA | Yes | Yes | Yes | Yes | Yes | |
| Gentamicin | Basal | 4 | cysS \|\| fusA | fusA | Yes | No | No | No | No | |
| Gentamicin | Basal | 5 | metQ \|\| ygfZ | ygfZ | Yes | No | No | No | No | |
| Gentamicin | Basal | 6 | fusA | fusA | Yes | No | No | No | No | |
| Gentamicin | Bile | 1 | ribB/yqiC \|\| ribB/yqiC | rib | Yes | Yes | Yes | Yes | Yes | |
| Gentamicin | Bile | 2 | | | No | No | No | No | No | No mutation identified |
| Gentamicin | Bile | 3 | dxs \|\| tfaP | dxs | Yes | No | No | No | No | |
| Gentamicin | Bile | 4 | ubiH | ubi | Yes | Yes | Yes | Yes | Yes | |
| Gentamicin | Bile | 5 | hemC | hem | Yes | Yes | Yes | Yes | Yes | |
| Gentamicin | Bile | 6 | ubiJ | ubi | Yes | No | No | No | Yes | |
| Gentamicin | pH | 1 | cpxA | cpxA | Yes | Yes | Yes | Yes | Yes | |
| Gentamicin | pH | 2 | ribB/yqiC | rib | No | No | No | No | No | Duplicated with GEN1Basal |
| Gentamicin | pH | 3 | ubiH | ubi | Yes | No | No | No | No | |
| Gentamicin | pH | 4 | ptsI \|\| nuoG | nuoG | Yes | No | No | No | Yes | |
| Gentamicin | pH | 5 | cpxA | cpxA | No | No | No | No | No | Duplicated with GEN2Basal |
| Gentamicin | pH | 6 | tufB \|\| hemB | tufB\|\|hem | Yes | No | No | Yes | Yes | Ambiguous family, grouped separately |

| | | | | | | | | | | |
|---|---|---|---|---|---|---|---|---|---|---|
| Gentamicin | Temperature | 1 | ribB/yqiC \|\| yncE/ansP | rib | Yes | Yes | Yes | Yes | Yes | |
| Gentamicin | Temperature | 2 | sbmA \|\| ribB/yqiC \|\| ribB/yqiC | rib | No | No | No | No | No | Duplicated with GEN1Basal |
| Gentamicin | Temperature | 3 | | | No | No | No | No | No | Not genotyped (Unable to culture) |
| Gentamicin | Temperature | 4 | ubiH | ubi | Yes | Yes | Yes | Yes | Yes | |
| Gentamicin | Temperature | 5 | hemL | hem | Yes | No | No | No | No | |
| Gentamicin | Temperature | 6 | ribE | rib | Yes | Yes | Yes | Yes | Yes | |
| Streptomycin | Basal | 1 | ribF | rib | Yes | Yes | Yes | Yes | Yes | |
| Streptomycin | Basal | 2 | dxs | dxs | Yes | No | No | No | No | |
| Streptomycin | Basal | 3 | ubiF | ubi | Yes | Yes | Yes | Yes | Yes | |
| Streptomycin | Basal | 4 | ribB/yqiC \|\| ribB/yqiC \|\| gcl | rib | No | No | No | No | No | Duplicated with STR5Basal |
| Streptomycin | Basal | 5 | yebT \|\| ribB/yqiC \|\| ribB/yqiC | rib | Yes | No | No | No | No | |
| Streptomycin | Basal | 6 | ribB/yqiC \|\| ribB/yqiC | rib | Yes | Yes | Yes | Yes | Yes | |
| Streptomycin | Bile | 1 | atpG | atpG | Yes | Yes | Yes | Yes | Yes | |
| Streptomycin | Bile | 2 | mngB/cydA | cydA | Yes | No | No | No | No | |
| Streptomycin | Bile | 3 | ubiF | ubi | Yes | No | No | No | No | |
| Streptomycin | Bile | 4 | ribD | rib | Yes | No | No | No | No | |
| Streptomycin | Bile | 5 | ubiB | ubi | Yes | Yes | Yes | Yes | Yes | |
| Streptomycin | Bile | 6 | ribE | rib | Yes | No | No | No | No | |
| Streptomycin | pH | 1 | rpsL \|\| arcB | rpsL | Yes | Yes | Yes | Yes | Yes | |
| Streptomycin | pH | 2 | rsmG | rsmG | Yes | Yes | Yes | Yes | Yes | |
| Streptomycin | pH | 3 | rpsL | rpsL | Yes | No | No | No | No | |

| | | | | | | | | | | |
|---|---|---|---|---|---|---|---|---|---|---|
| Streptomycin | pH | 4 | arcB \|\| rpsL | rpsL | Yes | Yes | Yes | Yes | Yes | |
| Streptomycin | pH | 5 | tufB | tufB | Yes | Yes | Yes | Yes | Yes | |
| Streptomycin | pH | 6 | ykg-ddlA deletion | hem | Yes | No | No | No | No | Δ83kb (78 genes) including hemB |
| Streptomycin | Temperature | 1 | atpG | atpG | Yes | Yes | Yes | Yes | Yes | |
| Streptomycin | Temperature | 2 | | | No | No | No | No | No | Not genotyped (Unable to culture) |
| Trimethoprim | Basal | 1 | djlA/yabP | djlA/yabP | Yes | Yes | Yes | Yes | Yes | |
| Trimethoprim | Basal | 2 | folA | folA | Yes | No | Yes | Yes | Yes | |
| Trimethoprim | Basal | 3 | phoP | pho | Yes | Yes | Yes | Yes | Yes | |
| Trimethoprim | Basal | 4 | folA | folA | No | No | No | No | No | Duplicated with TRM2Basal |
| Trimethoprim | Basal | 5 | kefC/folA | folA | Yes | No | Yes | Yes | Yes | |
| Trimethoprim | Basal | 6 | folA | folA | No | No | No | No | No | Duplicated with TRM6pH |
| Trimethoprim | Bile | 1 | folA | folA | Yes | Yes | Yes | Yes | Yes | |
| Trimethoprim | Bile | 2 | phoP | pho | No | No | No | No | No | Duplicated with TRM3Basal |
| Trimethoprim | Bile | 3 | mgrB | mgrB | Yes | Yes | Yes | Yes | Yes | |
| Trimethoprim | Bile | 4 | phoQ | pho | Yes | No | No | Yes | Yes | |
| Trimethoprim | Bile | 5 | mgrB/yobH | mgrB | Yes | No | No | No | No | |
| Trimethoprim | Bile | 6 | ybhH \|\| phoP | pho | Yes | Yes | Yes | Yes | Yes | |
| Trimethoprim | pH | 1 | | | No | No | No | No | No | Not genotyped (Unable to culture) |
| Trimethoprim | pH | 2 | gtrS | gtrS | Yes | Yes | Yes | Yes | Yes | |
| Trimethoprim | pH | 3 | folA | folA | Yes | Yes | Yes | Yes | Yes | |
| Trimethoprim | pH | 4 | phoQ | pho | No | No | No | No | No | Duplicated with TRM1Temp |
| Trimethoprim | pH | 5 | phoP \|\| fucP | pho | Yes | Yes | Yes | Yes | Yes | |

| | | | | | | | | | | |
|---|---|---|---|---|---|---|---|---|---|---|
| Trimethoprim | pH | 6 | folA | folA | Yes | No | Yes | Yes | Yes | |
| Trimethoprim | Temperature | 1 | phoQ | pho | Yes | Yes | Yes | Yes | Yes | |
| Trimethoprim | Temperature | 2 | phoQ | pho | Yes | Yes | Yes | Yes | Yes | |
| Trimethoprim | Temperature | 3 | phoQ \|\| alaV/rrlH | pho | Yes | Yes | Yes | Yes | Yes | |
| Trimethoprim | Temperature | 4 | folA | folA | No | No | No | No | No | Duplicated with TRM2Basal |
| Trimethoprim | Temperature | 5 | phoQ \|\| uacT | pho | Yes | Yes | Yes | Yes | Yes | |
| Trimethoprim | Temperature | 6 | phoP | pho | Yes | Yes | Yes | Yes | Yes | |
| Totals | | 113 | 95 | | 85 | 61 | 64 | 66 | 68 | |

**Table S1: Mutants used in this study.** This table shows all 113 mutants isolated in the resistance selection screen. The first three columns show the antibiotic used for selection, the environment where selection occurred, which replicate isolate (for the treatment) the mutant corresponds to, giving 113 independent mutants. After genotyping we identified mutations at high frequency (see methods) for 95 mutants with full genotype giving all genes with where mutations were at high frequency where a forward slash (/) between two genes means that the mutation occurred between the two genes and horizontal bars (||) separate different mutations that were found at high frequency in a mutant. We then grouped these full genotypes into mutants where genes in the same family were mutated, in order to better model the effects of genotype. When assigning a gene family for these mutants we used genes mutated in multiple independent lines (e.g. all genotypes with where rib mutations are grouped together rather than multiple different unrelated groups according other mutated genes). Of the genotyped mutants we measured resistance phenotypes for 85 mutants, excluding those where other mutants had very similar genotypes. We were able to use between 61-68 of these isolates for models of $IC_{90}$ to selection drug, growth at selection concentration (GASC), $IC_{90}$ to the paired drugs and antibiotic free growth (having to exclude strains with poor replication, see methods). The rightmost column gives additional information, particularly why mutants were not phenotyped. Here shorthand is used to describe mutants (e.g. TRM1Temp is the first replicate strain selected with trimethoprim at high temperature).

| Selection Drug | Assay Drug | Phenotype | Model A : Genotype | | | | | Model B: Selection Conditions | | | | |
|---|---|---|---|---|---|---|---|---|---|---|---|---|
| | | | Random Effects | | Fixed Effects | | | Random Effects | | Fixed Effects | | |
| | | | Strain | Strain : Block | Assay Environment | Genotype | Interaction | Strain | Strain : Block | Assay Environment | Selection Environment | Sympatry |
| Cefuroxime | Cefuroxime | $IC_{90}$ | ■ | ■ | ■ | ■ | ■ | ■ | ■ | ■ | | |
| Chloramphenicol | Chloramphenicol | $IC_{90}$ | | ■ | ■ | ■ | ■ | | ■ | ■ | | |
| Gentamicin | Gentamicin | $IC_{90}$ | | | ■ | ■ | | ■ | | ■ | | |
| Streptomycin | Streptomycin | $IC_{90}$ | ■ | ■ | ■ | ■ | ■ | ■ | | ■ | | |
| Trimethoprim | Trimethoprim | $IC_{90}$ | | ■ | | ■ | | ■ | | | | |
| Cefuroxime | Cefuroxime | GASC | ■ | ■ | ■ | ■ | ■ | ■ | ■ | | ■ | |
| Chloramphenicol | Chloramphenicol | GASC | | ■ | ■ | ■ | ■ | | | ■ | | |
| Gentamicin | Gentamicin | GASC | | | ■ | ■ | | | | ■ | | |
| Streptomycin | Streptomycin | GASC | | | ■ | ■ | ■ | ■ | | | | |
| Trimethoprim | Trimethoprim | GASC | ■ | ■ | ■ | ■ | ■ | ■ | ■ | ■ | | ■ |
| Cefuroxime | Gentamicin | $IC_{90}$ | ■ | ■ | ■ | ■ | ■ | ■ | ■ | ■ | | |
| Chloramphenicol | Polymyxin B | $IC_{90}$ | | ■ | ■ | ■ | ■ | | ■ | ■ | | |
| Gentamicin | Cefuroxime | $IC_{90}$ | | | ■ | ■ | ■ | ■ | | | | |
| Streptomycin | Tetracycline | $IC_{90}$ | | ■ | | | | | ■ | | ■ | |
| Trimethoprim | Nitrofurantoin | $IC_{90}$ | | ■ | ■ | ■ | | | ■ | ■ | ■ | |
| Cefuroxime | None | Cost | ■ | ■ | ■ | ■ | ■ | ■ | ■ | ■ | | |
| Chloramphenicol | None | Cost | | ■ | ■ | ■ | ■ | ■ | ■ | ■ | | |
| Gentamicin | None | Cost | ■ | ■ | ■ | ■ | ■ | ■ | ■ | | | |
| Streptomycin | None | Cost | ■ | ■ | ■ | ■ | ■ | ■ | ■ | ■ | | |
| Trimethoprim | None | Cost | ■ | ■ | ■ | ■ | ■ | ■ | ■ | ■ | ■ | |

**Table S2: Terms included in the minimal models reported in the text (black cells).** Models A and B were separately fitted to the different phenotypes. The models were independently simplified to the minimal models by dropping non-significant terms (not-included in higher order interactions), as described in more detail in the methods.
